# Rapid induction of antigen-specific CD4^+^ T cells guides coordinated humoral and cellular immune responses to SARS-CoV-2 mRNA vaccination

**DOI:** 10.1101/2021.04.21.440862

**Authors:** Mark M. Painter, Divij Mathew, Rishi R. Goel, Sokratis A. Apostolidis, Ajinkya Pattekar, Oliva Kuthuru, Amy E. Baxter, Ramin S. Herati, Derek A. Oldridge, Sigrid Gouma, Philip Hicks, Sarah Dysinger, Kendall A. Lundgreen, Leticia Kuri-Cervantes, Sharon Adamski, Amanda Hicks, Scott Korte, Josephine R. Giles, Madison E. Weirick, Christopher M. McAllister, Jeanette Dougherty, Sherea Long, Kurt D’Andrea, Jacob T. Hamilton, Michael R. Betts, Paul Bates, Scott E. Hensley, Alba Grifoni, Daniela Weiskopf, Alessandro Sette, Allison R. Greenplate, E. John Wherry

## Abstract

The SARS-CoV-2 mRNA vaccines have shown remarkable clinical efficacy, but questions remain about the nature and kinetics of T cell priming. We performed longitudinal antigen-specific T cell analyses in healthy individuals following mRNA vaccination. Vaccination induced rapid near-maximal antigen-specific CD4^+^ T cell responses in all subjects after the first vaccine dose. CD8^+^ T cell responses developed gradually after the first and second dose and were variable. Vaccine-induced T cells had central memory characteristics and included both Tfh and Th1 subsets, similar to natural infection. Th1 and Tfh responses following the first dose predicted post-boost CD8^+^ T cell and neutralizing antibody levels, respectively. Integrated analysis of 26 antigen-specific T cell and humoral responses revealed coordinated features of the immune response to vaccination. Lastly, whereas booster vaccination improved CD4^+^ and CD8^+^ T cell responses in SARS-CoV-2 naïve subjects, the second vaccine dose had little effect on T cell responses in SARS-CoV-2 recovered individuals. Thus, longitudinal analysis revealed robust T cell responses to mRNA vaccination and highlighted early induction of antigen-specific CD4^+^ T cells.

**Graphical Abstract:** 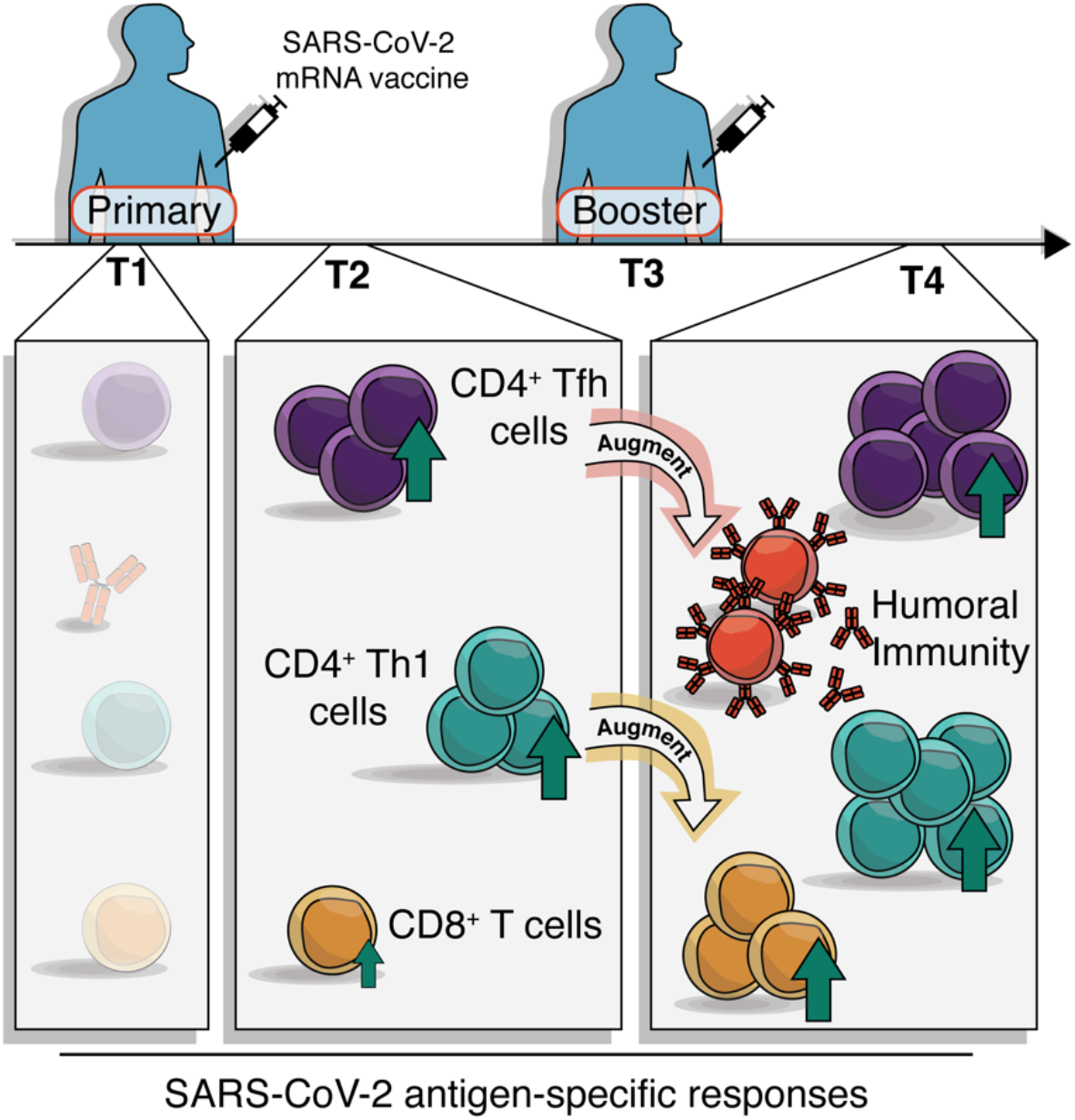

## Introduction

The COVID-19 pandemic has had a profound global toll on human life and socioeconomic well-being, prompting emergency use authorization of prophylactic mRNA vaccines (Cutler and Summers, 2020). Recent studies have documented strong antibody and memory B cell responses post-vaccination that neutralize SARS-CoV-2, including variants of concern (VOC) such as B.1.351 (Goel et al., 2021; Krammer et al., 2021; Sahin et al., 2020; Widge et al., 2021). B cells and antibodies are important components of immunological memory and antibody responses are the surrogate of protection for most licensed vaccines. However, patients who failed to develop neutralizing antibodies, in some cases due to inherited or treatment-induced B cell deficiencies, have recovered from COVID-19 (Soresina et al., 2020; Wu et al., 2020; Wurm et al., 2020). Moreover, in patients with hematological malignancy, CD8^+^ T cells appear to compensate for lack of humoral immunity and were associated with improved outcomes, indicating a role for T cells in protection against SARS-CoV-2 infection (Huang et al., 2021). The T cell response to mRNA vaccination is less well-characterized than the humoral response, though initial reports indicate that T cells, particularly CD4^+^ T cells, are primed by the vaccine (Anderson et al., 2020; Angyal, 2021; Camara et al., 2021; Jackson et al., 2020; Kalimuddin et al., 2021; Lederer et al., 2020; Mazzoni et al., 2021; Prendecki et al., 2021; Sahin et al., 2020; Stamatatos et al., 2021; Tarke et al., 2021b; Woldemeskel et al., 2021). However, the details of antigen-specific T cell induction following vaccination remain incompletely understood, and questions remain about the trajectory of the adaptive immune response following vaccination.

T cell immunity is functionally heterogeneous, with subsets of both CD4^+^ and CD8^+^ T cells contributing to protective immunity and long-term immunological memory. Specifically, CD4^+^ T follicular helper (Tfh) cells have key roles in the development of memory B cells, plasma cells and antibodies, whereas Th1 cells support and enhance the quality of memory CD8^+^ T cell responses (Crotty, 2011; Krawczyk et al., 2007; Luckheeram et al., 2012; Williams et al., 2006). In addition, the central memory or effector memory differentiation states of CD4^+^ and CD8^+^ T cells have implications for durability, recirculation, tissue access and responses upon antigen re-exposure (Kaech et al., 2002). In the context of mRNA vaccination, relatively little is known about the nature and differentiation state of antigen-specific CD4^+^ and CD8^+^ T cells. For example, it is unclear whether Tfh cells are efficiently primed and whether these cells relate to vaccine induced antibodies or memory B cells. It is also unclear whether the kinetics of T cell priming differs for CD4^+^ and CD8^+^ T cells, and how such T cell priming events might differ for SARS-CoV-2 naïve versus recovered subjects. Overall, the orchestration of different vaccine induced immune responses remains to be fully understood.

In this study we sought to address these questions and define the kinetics and differentiation state of vaccine-induced CD4^+^ and CD8^+^ T cells following mRNA vaccination. Nearly all SARS-CoV-2 naïve subjects mounted robust CD4^+^ T cell responses following the first vaccine dose, and the second dose further boosted both CD4^+^ and CD8^+^ T cell responses. In contrast, SARS-CoV-2 recovered individuals had maximal CD4^+^ and CD8^+^ T cell responses following the first dose of mRNA vaccine, and there was little additional T cell boosting after the second dose. Both CD4^+^ and CD8^+^ T cell responses were dominated by central memory-like cells, similar to memory T cells generated following natural infection. For CD4^+^ T cells, both Tfh and Th1 responses were efficiently generated following primary vaccination and strongly correlated with post-boost neutralizing antibody and CD8^+^ T cell responses, respectively. Finally, integrated analysis of 26 individual measures of antigen-specific T and B cells revealed coordinated immune response patterns and provided a comprehensive assessment of how antigen-specific adaptive immunity is shaped by mRNA vaccination.

## Results

We acquired longitudinal peripheral blood samples from a cohort of 29 SARS-CoV-2 naive and 10 SARS-CoV-2 recovered individuals who received mRNA vaccines through the University of Pennsylvania Health System (**Table S1**). We obtained peripheral blood mononuclear cells (PBMCs) at 4 key timepoints (**Fig. 1A**): pre-vaccine baseline (timepoint 1), two weeks post-primary vaccination (timepoint 2), the day of the booster vaccination (timepoint 3), and one week post-boost (timepoint 4). PBMCs from each of these timepoints were stimulated with peptide megapools containing SARS-CoV-2 spike epitopes optimized for presentation by MHC-I (CD8-E) or MHC-II (CD4-S) (Grifoni et al., 2020b; Tarke et al., 2021a). We then assessed peptide-dependent activation induced marker (AIM) expression by flow cytometry compared to unstimulated control samples (**Fig. 1A and Fig. S1A**) (Betts et al., 2003; Reiss et al., 2017). AIM^+^ CD4^+^ T cells were defined by dual-expression of CD200 and CD40L. Although dual expression of IFN-γ and 41BB was useful to visualize AIM^+^ CD8^+^ T cell populations (**Fig. 1A**), a two-marker strategy alone was sub-optimal for detecting vaccine-elicited responses due to high baseline signals (**Fig. S1B**). Thus, AIM^+^ CD8^+^ T cells were defined by expression of at least four of five markers: CD200, CD40L, 41BB, CD107a, and intracellular IFN-γ (**Fig. S1C**).

**Fig. 1:**
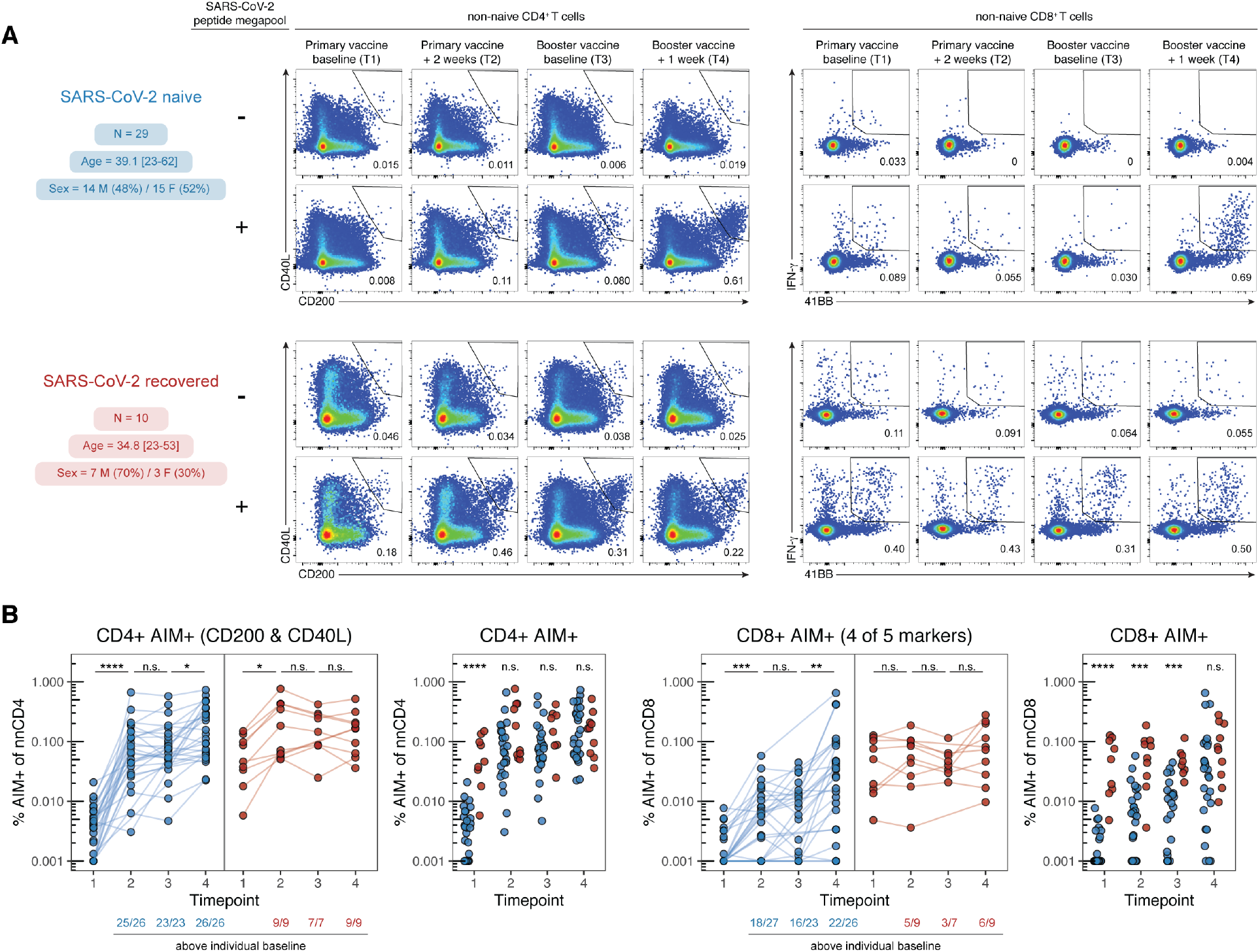
mRNA vaccination elicits antigen-specific CD4^+^ and CD8^+^ T cell responses. (A) Longitudinal study design and representative flow cytometry plots for identifying AIM^+^ CD4^+^ T cells (left) and visualizing AIM^+^ CD8^+^ T cells (right). Numbers represent the frequency of total non-naïve CD4^+^ or CD8^+^ T cells. (B) Summary plots of AIM^+^ CD4^+^ (left) and CD8^+^ (right) T cells defined as indicated above each plot. Values represent the frequency of AIM^+^ non-naïve cells after subtracting the frequency from paired unstimulated samples. Solid lines connect individual donors sampled longitudinally. Statistics were calculated using unpaired Wilcoxon test. Blue indicates SARS-CoV-2 naïve, red indicates SARS-CoV-2 recovered individuals.

As expected, most SARS-CoV-2 recovered donors had clearly detectable antigen-specific CD4^+^ and CD8^+^ T cell populations at baseline (**Fig. 1B**). In contrast, pre-vaccination responses to peptide stimulation were mostly undetectable in SARS-CoV-2 naïve individuals, though low levels of pre-vaccination AIM^+^ T cells were observed in some subjects that may be attributed to cross-reactive cells from a prior seasonal coronavirus infection (Grifoni et al., 2020a) (**Fig. 1B**). SARS-CoV-2 spike-specific CD4^+^ T cells were robustly primed in SARS-CoV-2 naïve and recovered individuals following the first dose of mRNA vaccine (**Fig. 1B**). SARS-CoV-2 naïve individuals, but not recovered individuals, received an additional boost to antigen-specific CD4^+^ T cells following the second vaccine dose (**Fig. 1B**). Overall, mRNA vaccination induced a universal CD4^+^ T cell response, as all individuals, regardless of prior infection with SARS-CoV-2, had greater frequencies of AIM^+^ CD4^+^ T cells post-boost than at baseline (**Fig. S1D**).

In contrast to the rapid and universal induction of spike-specific CD4^+^ T cells, SARS-CoV-2-specific CD8^+^ T cell responses developed more gradually and with greater variability in naïve individuals. Only 18 of 27 (67%) of SARS-CoV-2 naïve subjects generated a detectable antigen-specific CD8^+^ T cell response following the first dose. These CD8^+^ T cell responses were boosted by the second dose, and though the magnitude of response was variable, 22 of 26 SARS-CoV-2 naïve individuals (85%) had post-boost CD8^+^ T cell responses detectable above their individual pre-vaccine baseline (**Fig. 1B and Fig. S1D**). Individuals who had previously recovered from SARS-CoV-2 infection experienced no significant increase in the frequency of AIM^+^ CD8^+^ T cells from either dose of vaccine (**Fig. 1B and Fig. S1D**). A subset of recovered individuals (67%) did appear to have increased AIM^+^ CD8^+^ T cell frequencies compared to baseline, but as a group this increase did not reach statistical significance (**Fig. S1D**). In contrast to the modestly weaker induction of antibodies and memory B cells with increasing age observed in this cohort and others (Abu Jabal et al., 2021; Goel et al., 2021; Levi et al., 2021; Prendecki et al., 2021), T cell responses upon mRNA vaccination were not correlated with age (**Fig. S1E**). Taken together, these data demonstrate robust induction of antigen-specific T cell responses following mRNA vaccination, with more consistent induction of CD4^+^ T cell responses compared to CD8^+^ T cell responses.

We next sought to define the differentiation state of vaccine-induced AIM^+^ T cells. We first examined subsets of central and effector memory populations using CD45RA, CD27 and CCR7 (Hamann et al., 1997; Sallusto et al., 1999). With these markers, we defined central memory (CM), effector memory types 1, 2, and 3 (EM1, EM2, EM3) and terminally differentiated effector memory (EMRA) cells (**Fig. 2A, 2C, and S1A**) (Mathew et al., 2020). Total non-naïve CD4^+^ T cells were predominantly CM (CD45RA^−^ CD27^+^ CCR7^+^) in this cohort and the overall frequencies of these subsets were unchanged by vaccination (**Fig. S2A**). The baseline AIM^+^ CD4^+^ T cell response in SARS-CoV-2 recovered individuals, presumably generated during prior SARS-CoV-2 infection, was composed mainly of EM1 (CD45RA^−^ CD27^+^ CCR7^−^) and CM cells (**Fig. 2A-B**). The memory T cell subset distribution of these SARS-CoV-2 specific CD4^+^ T cells did not change dramatically following vaccination (**Fig. 2B**). In SARS-CoV-2 naïve individuals, the first dose of vaccine primarily induced AIM^+^ CD4^+^ T cells in the EM1 and CM subsets, similar to the response in recovered donors (**Fig. 2B**). Antigen-specific CD4^+^ EM2 (CD45RA^−^ CD27^−^ CCR7^+^) and EM3 (CD45RA^−^ CD27^−^ CCR7^−^) T cells, which share more effector-like properties (Romero et al., 2007), were also boosted by the vaccine, but remained minority populations compared to CM and EM1 (**Fig. 2B**).

**Fig. 2:**
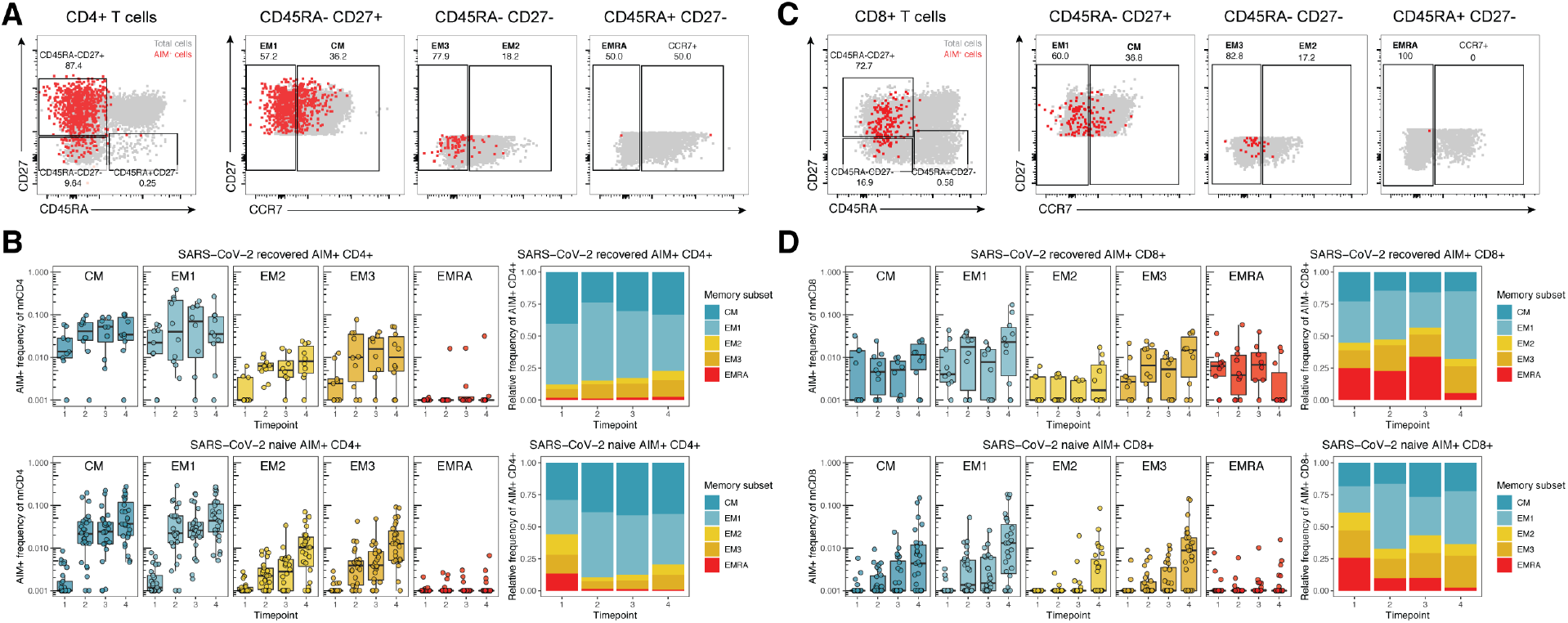
mRNA vaccination induces antigen-specific memory T cells that mirror memory T cell responses from natural infection. (A,C) Representative flow cytometric plots depicting the gating of AIM^+^ CD4^+^ (A) and CD8^+^ (C) T cells to identify the indicated memory T cell subsets in a SARS-CoV-2 naïve donor at timepoint 4. Red events depict AIM^+^ cells, gray events depict total CD4^+^ (A) or CD8^+^ (C) T cells from the same donor. Numbers indicate the frequency of AIM^+^ cells falling within each gate. (B,D) Frequency of memory T cell subsets in AIM^+^ CD4^+^ (B) and AIM^+^ CD8^+^ (D) T cells. Top panels depict SARS-CoV-2 recovered donors. Bottom panels depict SARS-CoV-2 naïve donors. Left panels depict the background-subtracted percent of non-naïve T cells that are AIM^+^ cells of each subset. Right panels depict the relative frequency of each memory T cell subset in the background-subtracted AIM^+^ population. CM = CD45RA^−^ CD27^+^ CCR7^+^, EM1 = CD45RA^−^ CD27^+^ CCR7^−^, EM2 = CD45RA^−^ CD27^−^ CCR7^+^, EM3 = CD45RA^−^ CD27^−^ CCR7^−^, EMRA = CD45RA^+^ CD27^−^ CCR7^−^. Timepoints are as defined in Fig. 1A.

AIM^+^ CD8^+^ T cells had a similar subset distribution to AIM^+^ CD4^+^ T cells. Total non-naïve CD8^+^ T cells were distributed throughout memory T cell subsets and the frequencies of these subsets were unchanged by vaccination (**Fig S2B**). The baseline antigen-specific CD8^+^ T cell response in recovered subjects was composed of similar proportions of EM1, CM, and terminally-differentiated CD8^+^ EMRA (CD45RA^+^ CD27^−^ CCR7^−^) T cells (**Fig. 2C-D**). A smaller proportion of AIM^+^ EM2 and EM3 CD8^+^ T cells was observed at baseline in recovered subjects. These proportions stayed relatively consistent throughout the course of vaccination in recovered subjects, and there were no statistically significant changes from baseline (**Fig. 2D**). In contrast, in SARS-CoV-2 naive individuals, few AIM^+^ EMRA CD8^+^ T cells were observed at any time point (**Fig. 2D**). Rather, vaccine-primed AIM^+^ CD8^+^ T cells in these subjects were largely EM1 with minority populations of CM and EM3 cells (**Fig. 2D**). With the exception of the EMRA population, the antigen-specific AIM^+^ CD8^+^ T cell response in SARS-CoV-2 naïve donors following vaccination resembled that observed in recovered donors (**Fig. 2D**). These data indicate that the vaccine-elicited T cell response has a similar memory T cell subset distribution to the response generated following SARS-CoV-2 infection and is comprised of primarily CD45RA^−^ CD27^+^ memory T cells.

Given the role of helper subsets like CD4^+^ T follicular helper cells (Tfh) to help B cell responses and the importance of Th1 cells in viral infections, we next explored the differentiation state of AIM^+^ CD4^+^ T cells. To this end, we examined antigen-specific CXCR5^+^ Tfh in circulation (cTfh) as well as CXCR5^−^ Th1 (CXCR3^+^CCR6^−^), Th2 (CXCR3^−^CCR6^−^), Th17 (CXCR3^−^CCR6^+^), and Th1/17 (CXCR3^+^CCR6^+^) cells (**Fig. 3A and S1A**) (Acosta-Rodriguez et al., 2007; Schmitt et al., 2014; Trifari et al., 2009). Total non-naive CD4^+^ T cell populations predominantly had Th1 and Th2 phenotypes (**Fig. S3A**). The baseline AIM^+^ CD4^+^ T cell response in recovered individuals, however, was dominated by cTfh and Th1 cells (**Fig. 3A-B**). The first dose of vaccine led to further expansion of AIM^+^ cTfh and Th1 cells in these recovered subjects, and this pattern was largely maintained through the course of vaccination (**Fig. 3B**). In SARS-CoV-2 naïve subjects, the first vaccine dose also elicited predominantly antigen-specific Th1 and cTfh cells (**Fig. 3B**). This distribution was sustained through booster vaccination in SARS-CoV-2 naïve individuals, with these AIM^+^ subsets being further boosted by the second vaccine dose (**Fig. 3B**). Thus, the vaccine-elicited AIM^+^ CD4^+^ T cell response to mRNA vaccination qualitatively resembled the response to natural infection and was characterized by robust induction of antigen-specific cTfh and Th1 cells.

**Fig. 3:**
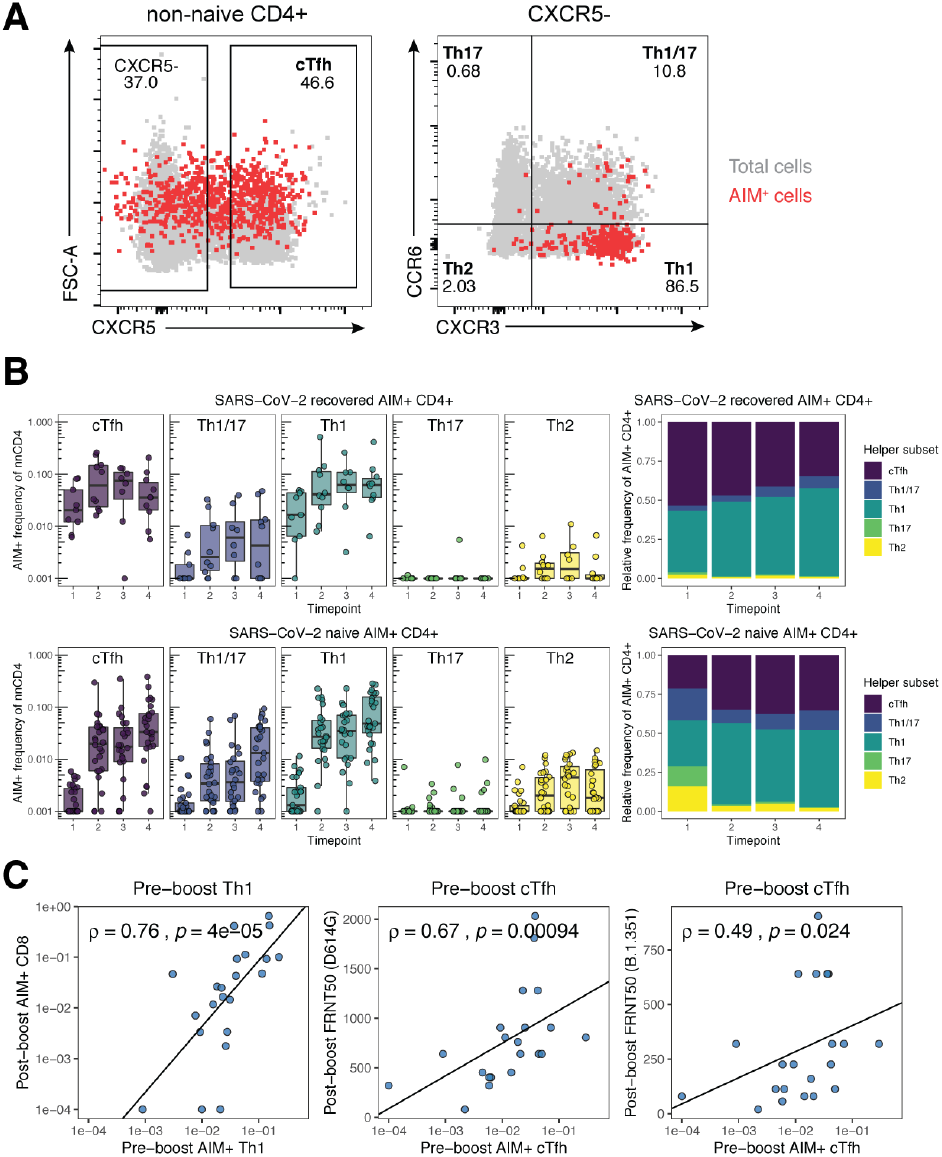
Early antigen-specific CD4^+^ helper T cell responses shape humoral and cellular adaptive immune responses to mRNA vaccination. (A) Representative flow cytometric plots depicting the gating of AIM^+^ CD4^+^ T cells to identify the indicated helper subsets in a SARS-CoV-2 naïve donor at timepoint 4. Red events depict AIM^+^ T cells, gray events depict total CD4^+^ T cells from the same donor. (B) Frequency of T helper subsets in AIM^+^ CD4^+^ T cells. Top panel depicts SARS-CoV-2 recovered donors. Bottom panel depicts SARS-CoV-2 naïve donors. Left panel depicts the background-subtracted percent of non-naïve CD4^+^ T cells that are AIM^+^ helper T cells in each subset. Right panel depicts the relative frequency of each helper T cell subset in the background-subtracted AIM^+^ population. cTfh = CXCR5^+^ of non-naïve CD4^+^ T cells, Th1 = CXCR5^−^ CXCR3^+^ CCR6^−^, Th2 = CXCR5^−^ CXCR3^−^ CCR6^−^, Th17 = CXCR5^−^ CXCR3^−^ CCR6^+^, Th1/17 = CXCR5^−^ CXCR3^+^ CCR6^+^. (C) Correlations between the frequency of pre-boost (timepoint 2) AIM^+^ Th1 or AIM^+^ cTfh cells with post-boost (timepoint 4) AIM^+^ CD8^+^ T cells or neutralizing titers against dominant (D614G) or variant (B.1.351) strains of SARS-CoV-2 as published in a previous study of the same cohort (Goel et al., 2021). FRNT50 = Focus reduction neutralization titer 50&. Only SARS-CoV-2 naïve donors were considered for these correlations. Associations were calculated using Spearman rank correlation and are shown with Pearson trend lines for visualization. Timepoints are as defined in Fig. 1A.

The rapid induction of antigen-specific CD4^+^ T cells following the first mRNA vaccine dose, particularly Th1 and cTfh cells, may provide a population of helper T cells available to enhance immune responses to the second vaccine dose. Th1 cells predominantly facilitate the CD8^+^ T cell response, whereas Tfh cells help foster optimal B cell, germinal center, and antibody responses (Crotty, 2011; Krawczyk et al., 2007; Luckheeram et al., 2012; Williams et al., 2006). Indeed, we observed a strong correlation between the frequency of pre-boost AIM^+^ Th1 cells and the frequency of post-boost AIM^+^ CD8^+^ T cells in SARS-CoV-2 naïve individuals (**Fig. 3C**), consistent with a role for Th1 cells generated by primary vaccination in enhancing the CD8^+^ T cell responses following booster vaccination. Similarly, the frequency of pre-boost antigen-specific cTfh cells correlated with post-boost neutralizing antibody titers against both the dominant strain of SARS-CoV-2 (D614G, dominant at the time of study) and the B.1.351 variant (Goel et al., 2021) (**Fig. 3C**). Pre-boost Th1 did not significantly correlate with post-boost neutralizing titers, nor did pre-boost cTfh correlate with post-boost CD8^+^ T cell responses, supporting the distinct contributions of these pre-boost immune cell types to post-boost vaccine-elicited immune responses (**Fig. S3B**). Moreover, baseline AIM^+^ Th1 and cTfh cells in SARS-CoV-2 naïve subjects did not correlate with post-boost CD8^+^ T cell or neutralizing responses, respectively, suggesting minimal contribution of pre-existing cross-reactive CD4^+^ T cells to the observed immune response to SARS-CoV-2 mRNA vaccines (**Fig. S3C**). These observations highlight a key functional role for vaccine-elicited CD4^+^ T cells and suggest possible downstream effects of skewed antigen-specific CD4^+^ T cell responses. Moreover, these data highlight one of the potential benefits of a two-dose vaccination regimen, where CD4^+^ T cells primed by the first vaccine dose may augment and coordinate responses following the booster vaccination.

These CD4^+^ T cell data suggested important interrelationships between distinct immune responses generated by mRNA vaccination. To further examine this notion of coordinated immune responses following vaccination, we compiled the antigen-specific T cell data described above with a previously reported dataset of antibody and memory B cell responses from this cohort (Goel et al., 2021). Using these data, we integrated 26 antigen-specific features of the immune response to mRNA vaccination into high-dimensional UMAP space (**Fig. 4A**). Correlating individual antigen-specific features with the UMAP coordinates revealed that UMAP1 is a measure of the anti-SARS-CoV-2 immune response to vaccination (**Fig. 4A-B**). UMAP1 also revealed a signal of previous SARS-CoV-2 infection, as recovered subjects occupied a location with increased UMAP1 signal at baseline (**Fig. 4A-B**). Specifically, UMAP1 captured a coordinated immune response in which antigen-specific CD8^+^ T cells and CD4^+^ Th1 and cTfh cells were increased coordinately with antibodies, IgG^+^ memory B cells, RBD-focused humoral responses, and increased neutralizing antibody titers (**Fig. 4D-E**). Total non-naive lymphocyte populations were not altered and did not correlate with the antigen-specific responses, consistent with induction of a targeted vaccine-elicited response (**Fig. S4A**). This UMAP projection also revealed trajectory shifts that were notable between naïve and recovered subjects. For example, in SARS-CoV-2 recovered individuals, there was an increase in both UMAP1 and UMAP2 following primary vaccination, but essentially no change following the second vaccine dose (**Fig. 4A-C**). In SARS-CoV-2 naïve subjects there was a more dynamic trajectory over time with an initial increase in UMAP1 and decrease in UMAP2 signal at timepoints 2 and 3, followed by a coalescence towards increased UMAP1 and UMAP2 after the second vaccine dose (**Fig. 4A-C**). UMAP2 captures the different kinetics of T cell and humoral responses in SARS-CoV-2 naïve individuals. A rapid antigen-specific CD4^+^ T cell, and to a lesser extent CD8^+^ T cell response drives the UMAP2 coordinate in a negative direction pre-boost, whereas robust humoral immunity post-boost drives UMAP2 in a positive direction (**Fig. 4C-E**). Finally, correlations of key antigen-specific parameters of the vaccine response over time revealed relationships between post-primary and post-boost immunity within and between arms of the adaptive immune system, highlighting pre-boost features like cTfh and Th1 that correlate with post-boost humoral and cellular responses (**Fig. S4B**). In summary, this unbiased integrated analysis of 26 antigen-specific immune responses illustrates the coordinated immunological underpinnings of the immunity induced by mRNA vaccines.

**Fig. 4:**
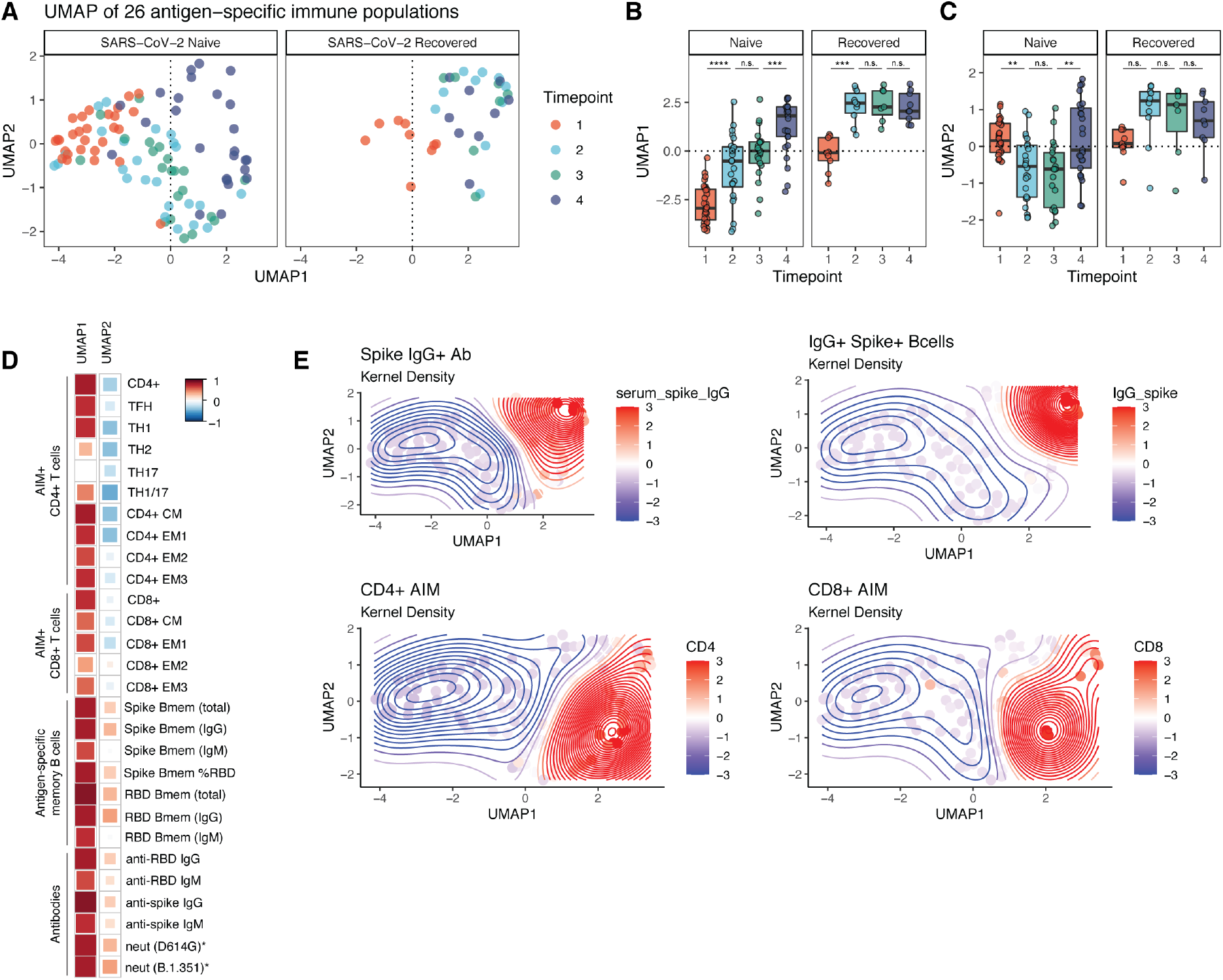
mRNA vaccination provokes a coordinated immune response in SARS-CoV-2 naive and recovered individuals. (A) UMAP projections of aggregated antigen-specific data for T cell, memory B cell, and antibody responses over time. Memory B cell and antibody data were taken from a previously-published dataset using the same cohort (Goel et al., 2021). Colors represent timepoints at which PBMCs were collected throughout the study. Parameters were considered as frequency of non-naïve T cells or memory B cells, capturing both the magnitude and skewing of responses. (B-C) Summary plots of UMAP1 (B) and UMAP2 (C) coordinates over time. Individual points represent individual participants. Statistics were calculated using unpaired Wilcoxon test. (D) Correlations of the individual antigen specific features used to train the UMAP against the UMAP1 and UMAP2 axis. (*) represents correlation of features against UMAP1 and UMAP2 that were not used to train the original UMAP. Red indicates positive correlations and blue indicates negative correlations. (E) Kernel density plots displaying the variation of selected antigen-specific features across UMAP space. Timepoints are as defined in Fig. 1A.

## Discussion

In this study, we interrogated the antigen-specific CD4^+^ and CD8^+^ T cell responses induced by SARS-CoV-2 mRNA vaccination in a longitudinal cohort of SARS-CoV-2 naïve and recovered individuals. Our data demonstrate robust induction of antigen-specific T cells by mRNA vaccination that may contribute, in addition to previously defined humoral responses, to durable protective immunity. In particular, antigen-specific memory CD4^+^ and CD8^+^ T cells are likely to be less impacted by antibody escape mutations in variant viral strains, as T cells can recognize peptide epitopes distributed throughout the SARS-CoV-2 spike protein (Angyal, 2021; Tarke et al., 2021b; Woldemeskel et al., 2021). Moreover, unlike vaccine-induced B cell and antibody responses, which have been noted to decrease with age (Abu Jabal et al., 2021; Goel et al., 2021; Levi et al., 2021; Prendecki et al., 2021), substantial age-associated changes in the induction of antigen-specific T cell responses were not observed. Finally, the generation of robust T cell responses by mRNA vaccines may have implications for long-term protective immunity, as memory CD4^+^ and CD8^+^ T cells can be exceptionally durable in other vaccine settings (Akondy et al., 2017; Hammarlund et al., 2003).

Vaccine-induced CD4^+^ and CD8^+^ T cells specific for SARS-CoV-2 were qualitatively similar to baseline memory T cell responses generated following natural SARS-CoV-2 infection and mainly mapped to CM and EM1 memory T cell subsets. These two subsets share many functional and memory-like attributes, but differ in CCR7 expression. Since CCR7 promotes homing to secondary lymphoid tissues, EM1 may represent memory T cells that can survey blood and peripheral tissues, whereas CM can home efficiently to lymphoid tissues (Romero et al., 2007). These memory T cell subsets are longer-lived compared to effector T cells, and access to secondary lymphoid tissues may allow CM cells to contribute to recall responses upon booster vaccination or future infection. Although we await follow-up studies to directly interrogate longevity, the observed induction of memory T cell subsets with capacity for durability by mRNA vaccination supports the hypothesis that vaccine-induced CD4^+^ and CD8^+^ T cell responses will be long-lived and capable of contributing to future recall responses.

One key observation was the rapid and universal induction of SARS-CoV-2-specific CD4^+^ T cells following the first vaccine dose in SARS-CoV-2 naïve individuals. This observation may be noteworthy given the gradual development of antigen-specific CD8^+^ T cells observed here and the previous observations for humoral responses (Goel et al., 2021; Jackson et al., 2020), which only consistently reach maximal levels after the second vaccine dose. These data point to the early induction of antigen-specific CD4^+^ T cells as a possible contributor to the protection observed in clinical trials as early as two weeks after the first vaccine dose (Baden et al., 2020; Polack et al., 2020), when neutralizing antibody levels are still low in many individuals (Goel et al., 2021). Indeed, CD4^+^ T cells can prevent symptomatic SARS-CoV infection in animal models (Zhao et al., 2016), and the rapid induction of antigen-specific CD4^+^ T cells after only a single vaccine dose may explain the disconnect between low neutralizing responses and vaccine-induced protective immunity following the first dose.

The notion that early CD4^+^ T cell responses have a functional role in immunity is also supported by the correlation between pre-boost Th1 and cTfh cells with post-boost CD8^+^ T cell and neutralizing antibody responses, respectively. In addition to strongly correlating with post-boost neutralizing antibody titers, pre-boost antigen-specific cTfh cells also correlated with post-boost memory B cell responses. When examining multiple individual antigen-specific responses over time, pre-boost cTfh were better predictors of post-boost humoral responses than many pre-boost readouts of humoral immunity, pointing to the critical role of Tfh in coordinating humoral immunity. Likewise, pre-boost antigen-specific CD4^+^ T cells, and especially Th1 cells, were more strongly correlated with post-boost CD8^+^ T cell responses than were pre-boost CD8^+^ T cells. These findings suggest that the CD4^+^ T cell response generated by the first vaccine dose guides multiple arms of the adaptive immune response to booster vaccination and highlight the benefits of a prime-boost strategy to amplify a coordinated vaccine-induced immune response.

Previous studies have demonstrated that individuals who have recovered from SARS-CoV-2 infection achieve maximum antigen-specific humoral immune responses after only a single vaccine dose, raising the question of whether a second vaccine dose is necessary in these individuals (Angyal, 2021; Bradley et al., 2021; Camara et al., 2021; Goel et al., 2021; Mazzoni et al., 2021; Saadat et al., 2021; Samanovic et al., 2021; Stamatatos et al., 2021). Our current studies now provide information on both CD4^+^ and CD8^+^ T cell responses in naïve and recovered subjects and support the idea that the second dose of vaccine has minimal impact on the magnitude, memory phenotype, or helper subset distribution of antigen-specific CD4^+^ or CD8^+^ T cell responses in SARS-CoV-2 recovered subjects. Moreover, an integrated analysis of 26 antigen-specific features of the immune response to vaccination highlighted the immunological benefit of the first dose in recovered subjects while also illustrating the relative stability of the immune landscape in response to the second vaccine dose. In contrast, in SARS-CoV-2 naïve subjects, there was robust and dynamic change in the coordination and evolution of the antigen-specific immune response following the first as well as the second vaccine dose. These data point to the immunological benefit of two vaccine doses in SARS-CoV-2 naïve subjects and highlight the coordination between different arms of the adaptive immune response following mRNA vaccination. In concert with robust humoral immunity, the preferential induction of Th1, Tfh, and central memory-like T cells indicates that the vaccine-elicited immune response is specifically focused on the key hallmarks of long-term antiviral immunity that are likely to confer lasting protection against SARS-CoV-2 infection.

## Acknowledgements

We would like to thank the study participants for their generosity in making the study possible. We also thank Wenzhao Meng, Aaron M. Rosenfeld, Eline T. Luning Prak, and the members of the Wherry lab for helpful discussions and feedback. This work was supported by grants from the NIH AI105343, AI082630, AI108545, AI155577, AI149680 (to EJW), AI152236 (to PB), AI082630 (to RSH), HL143613 (to JRG), P30-AI0450080 (to ELP), T32 AR076951-01 (to SAA), T32 CA009140 (to JRG and DM), T32 AI055400 (to PH), U19AI082630 (to SEH and EJW), funding from the Allen Institute for Immunology (to SAA, EJW), Cancer Research Institute-Mark Foundation Fellowship (to JRG), Chen Family Research Fund (to SAA), the Parker Institute for Cancer Immunotherapy (to JRG, EJW), the Penn Center for Research on Coronavirus and Other Emerging Pathogens (to PB), the University of Pennsylvania Perelman School of Medicine COVID Fund (to RRG, EJW), the University of Pennsylvania Perelman School of Medicine 21^st^ Century Scholar Fund (to RRG), and a philanthropic gift from Jeffrey Lurie, Joel Embiid, Josh Harris, and David Blitzer (to SEH). Work in the Wherry lab is supported by the Parker Institute for Cancer Immunotherapy. This work was also supported by NIH contract Nr. 75N9301900065 (to DW, AS).

## Author contributions

MMP, DM, and EJW concieved the study. MMP, DM, RRG, and SAA carried out experiments. SAA and OK were invovled in clinical recruitment. ELP, DAO and JRG provided input on statistical analyses. DM, AB, and RSH contributed to the methodology. MMP, DM, RRG, SA, AH, SK, KD, and JTH processed peripheral blood samples and managed the sample database. RRG, OK, JD, and SL performed phlebotomy. AP, SG, PH, SD, KAL, LKC, WM, AB, MEW, CMM, AG, DW, and AS provided data and materials. ARG and EJW supervised the study. All authors participated in data analysis and interpretation. MMP, DM, and EJW wrote the manuscript.

## STAR METHODS

### RESOURCE AVAILABILITY

#### Lead contact

Further information and requests for resources and reagents should be directed to and will be fulfilled by the lead contact, E. John Wherry.

#### Materials availability

This study did not generate new unique reagents.

#### Data and code availability

Raw data files and reagents are available from the authors upon request.

### EXPERIMENTAL MODEL AND SUBJECT DETAILS

#### Human subjects

39 individuals (29 SARS-CoV-2 naïve, 10 SARS-CoV-2 recovered) provided informed consent and were enrolled in the study with approval from the University of Pennsylvania Institutional Review Board (IRB# 844642). All participants were otherwise healthy and did not report any history of chronic health conditions. Subjects were identified as SARS-CoV-2 naïve or recovered via combined self-reporting and laboratory evidence of a prior SARS-CoV-2 infection. All subjects received either Pfizer (BNT162b2) or Moderna (mRNA-1273) mRNA vaccines. Samples were collected at 4 timepoints: pre-vaccine baseline (timepoint 1), two weeks post-primary vaccination (timepoint 2), the day of the booster vaccination (timepoint 3), and one week post-boost (timepoint 4). Each study visit included collection of clinical questionnaire data and 80-100mL of peripheral blood. Full cohort and demographic information is provided in **Table S1**.

### METHOD DETAILS

#### Sample processing

Venous blood was collected into sodium heparin and EDTA tubes by standard phlebotomy. Blood tubes were centrifuged at 3000rpm for 15 minutes to separate plasma. Heparin and EDTA plasma were stored at −80°C for paired serological analyses. Remaining whole blood was diluted 1:1 with RPMI 1640 (Corning) supplemented with 1% fetal bovine serum (FBS), 2mM L-Glutamine, 100 U/mL Penicillin, and 100 μg/mL Streptomycin (R1 medium) and layered onto SEPMATE tubes (STEMCELL Technologies) containing lymphoprep gradient (STEMCELL Technologies). SEPMATE tubes were centrifuged at 1200g for 10 minutes and the PBMC fraction was collected into new tubes and washed with R1. PBMCs were then treated with ACK lysis buffer (Thermo Fisher) for 5 minutes to lyse red blood cells. Samples were washed again with R1, passed through a 70μm cell strainer, and cell counts were acquired with a Countess automated cell counter (Thermo Fisher). PBMCs were cryopreserved in 10% DMSO in FBS.

#### Activation induced marker(AIM) expression assay

PBMCs were thawed by warming frozen cryovials in a 37°C water bath and resuspending cells in 10mL of RPMI supplemented with 10% FBS, 2mM L-Glutamine, 100 U/mL Penicillin, and 100 μg/mL Streptomycin (R10). Cells were washed once in R10, counted using a Countess automated cell counter (Thermo Fisher), and resuspended in fresh R10 to a density of 5×10^6^ cells/mL. For each condition, duplicate wells containing 1×10^6^ cells in 200μL were plated in 96-well round-bottom plates and rested overnight in a humidifed incubator at 37°C, 5% CO_2_. After 16 hours, CD40 blocking antibody (0.5μg/mL final concentration) was added to cultures for 15 minutes prior to stimulation. Cells were then stimulated for 24 hours with costimulation (anti-human CD28/CD49d, BD Biosciences) and peptide megapools (CD4-S for all CD4^+^ T cell analyses, CD8-E for all CD8^+^ T cell analyses) at a final concentration of 1 μg/mL. Peptide megapools were prepared as previously described (Grifoni et al., 2020b; Tarke et al., 2021a). Matched unstimulated samples for each donor at each timepoint were treated with costimulation alone. 20 hours post-stimulation, antibodies targeting CXCR3, CCR7, CD40L, CD107a, CXCR5, and CCR6 were added to the culture along with monensin (GolgiStop, BD Biosciences) for a four-hour stain at 37°C. After four hours, duplicate wells were pooled and cells were washed in PBS supplemented with 2% FBS (FACS buffer). Cells were stained for 10 minutes at room temperature with Ghost Dye Violet 510 and Fc receptor blocking solution (Human TruStain FcX™, BioLegend) and washed once in FACS buffer. Surface staining for 30 minutes at room temperature was then performed with antibodies directed against CD4, CD8, CD45RA, CD27, CD3, CD69, CD40L, CD200, OX40, and 41BB in FACS buffer. Cells were washed once in FACS buffer, fixed and permeabilizied for 30 minutes at room temperature (eBioscience™ Foxp3 / Transcription Factor Fixation/Permeabilization Concentrate and Diluent), and washed once in 1X Permeabilization Buffer prior to staining for intracellular IFN-γ overnight at 4°C. Cells were then washed once and resuspended in 1% paraformaldehyde in PBS prior to data acquisition.

All data from AIM expression assays were background-subtracted using paired unstimulated control samples. For memory T cell and helper T cell subsets, the AIM+ background frequency of non-naïve T cells was subtracted independently for each subset. AIM^+^ cells were identified from non-naïve T cell populations. AIM^+^ CD4^+^ T cells were defined by dual-expression of CD200 and CD40L. AIM^+^ CD8^+^ T cells were defined by a boolean analysis identifying cells expressing at least four of five markers: CD200, CD40L, 41BB, CD107a, and intracellular IFN-γ.

#### Flow cytometry

Data were acquired on a BD Symphony A5 instrument. Standardized SPHERO rainbow beads (Spherotech) were used to track and adjust photomultiplier tube voltages over time. Compensation was performed using UltraComp eBeads (Thermo Fisher). Up to 2×10^6^ events were acquired per sample. Data were analyzed using FlowJo v10 (BD Bioscience). A full gating strategy for segregation of T cell subsets is shown in **Fig. S1A**.

#### Antibody and memory B cell responses

The dataset of antibody and memory B cell responses from the same cohort of individuals was published previously (Goel et al., 2021).

### QUANTIFICATION AND STATISTICAL ANALYSIS

#### Data visualization and statistics

All data were analyzed using custom scripts in R and visualized using RStudio. Boxplots represent median with interquartile range. The 26 parameters used to train the UMAP were scaled by column (z-score normalization) prior to generating UMAP coordinates. Statistical tests are indicated in the corresponding figure legends. All tests were performed two-sided with a nominal significance threshold of p < 0.05. In all cases of multiple comparisons, adjustment was performed using Holm correction. Unpaired tests were used for comparisons between timepoints, as some participants lacked samples from individual timepoints. * indicates p < 0.05, ** indicates p < 0.01, *** indicates p < 0.001, **** indicates p < 0.0001. Source code and data files are available upon request from the authors.

**KEY RESOURCES TABLE.**

| REAGENT or RESOURCE | SOURCE | IDENTIFIER |
| --- | --- | --- |
| **Antibodies** |  |  |
| BUV395 CD4 | BD Biosciences | Cat#563550 |
| BUV496 CD8 | BD Biosciences | Cat#612943 |
| BUV615 CD45RA | BD Biosciences | Cat#751555 |
| BUV737 CD27 | BD Biosciences | Cat#612829 |
| BUV805 CD3 | BD Biosciences | Cat#612896 |
| BV421 CXCR3 | Biolegend | Cat#353716 |
| BV650 CCR7 | Biolegend | Cat#353234 |
| BV605 CD69 | Biolegend | Cat#310938 |
| BV711 CD40L | Biolegend | Cat#310838 |
| BV785 CD107a | Biolegend | Cat#328644 |
| FITC IFNγ | Biolegend | Cat#502515 |
| PE CD200 | Biolegend | Cat#399804 |
| PECy7 OX40 | Biolegend | Cat#350012 |
| AF647 41BB | Biolegend | Cat#309810 |
| APC R700 CXCR5 | BD Biosciences | Cat#565191 |
| APCCy7 CCR6 | Biolegend | Cat#353432 |
| **Chemicals, peptides, and recombinant proteins** |  |  |
| CD4-S peptide megapool | Synthetic Biomolecules (aka A&A) | http://www.syntheticbiomolecules.com/ |
| CD8-E peptide megapool | Synthetic Biomolecules (aka A&A) | http://www.syntheticbiomolecules.com/ |
| **Other** |  |  |
| Ghost Dye Violet 510 | Tonbo | Cat#13-0870-T500 |
| GolgiStop (Containing Monensin) | BD Biosciences | Cat#51-2092K7 |
| CD40 Antibody, anti-human, pure-functional grade | Miltenyi Biotech | Cat#130-094-133 |
| Anti-Human CD28/CD49d Purified | BD Biosciences | Cat#347690 |
| Human TruStain FcX™ (Fc Receptor Blocking Solution) | Biolegend | Cat#422302 |
| Foxp3 / Transcription Factor Fixation/Permeabilization Concentrate and Diluent | eBioscience |  |

## Supplementary Information

**Table S1: Cohort demographics.**

**Fig. S1: Gating strategy and magnitude of AIM^+^ T cell responses.**

**Fig. S2: Summary of memory T cell subsets in total non-naïve T cells.**

**Fig. S3: CD4^+^ helper T cell subsets supplement.**

**Fig. S4: Integrated analysis supplement.**

