## Supplementary Information for "Rapid induction of antigen-specific CD4^+^ T cells guides coordinated humoral and cellular immune responses to SARS-CoV-2 mRNA vaccination"

### Table S1: Cohort demographics.

| | | SARS-CoV-2 Naïve | SARS-CoV-2 Recovered |
|---|---|---|---|
| **N** | Number of individuals | 29 (74.4%) | 10 (25.6%) |
| **Age** | Mean years | 39.1 | 34.8 |
| | 20-30 | 7 (24.1%) | 4 (40%) |
| | 30-40 | 9 (31.0%) | 3 (30%) |
| | 40-50 | 8 (27.6%) | 1 (10%) |
| | 50+ | 5 (17.2%) | 2 (20%) |
| **Sex** | Male | 14(48.2%) | 7 (70%) |
| | Female | 15 (51.7%) | 4 (40%) |
| **Race/Ethnicity** | White: Non-Hispanic/Latino | 17 (58.6%) | 7 (70%) |
| | White: Hispanic/Latino | 3 (10.3%) | 1 (10%) |
| | Asian | 6 (20.7%) | 2 (20%) |
| | Black | 2 (6.9%) | 0 (0% |
| | Native | 0 (0%) | 1 (10%) |
| | Other | 1 (3.4%) | 0 (0%) |
| **Vaccine Type** | Pfizer | 29 (100%) | 7 (70%) |
| | Moderna | 0 (0%) | 3 (30%) |

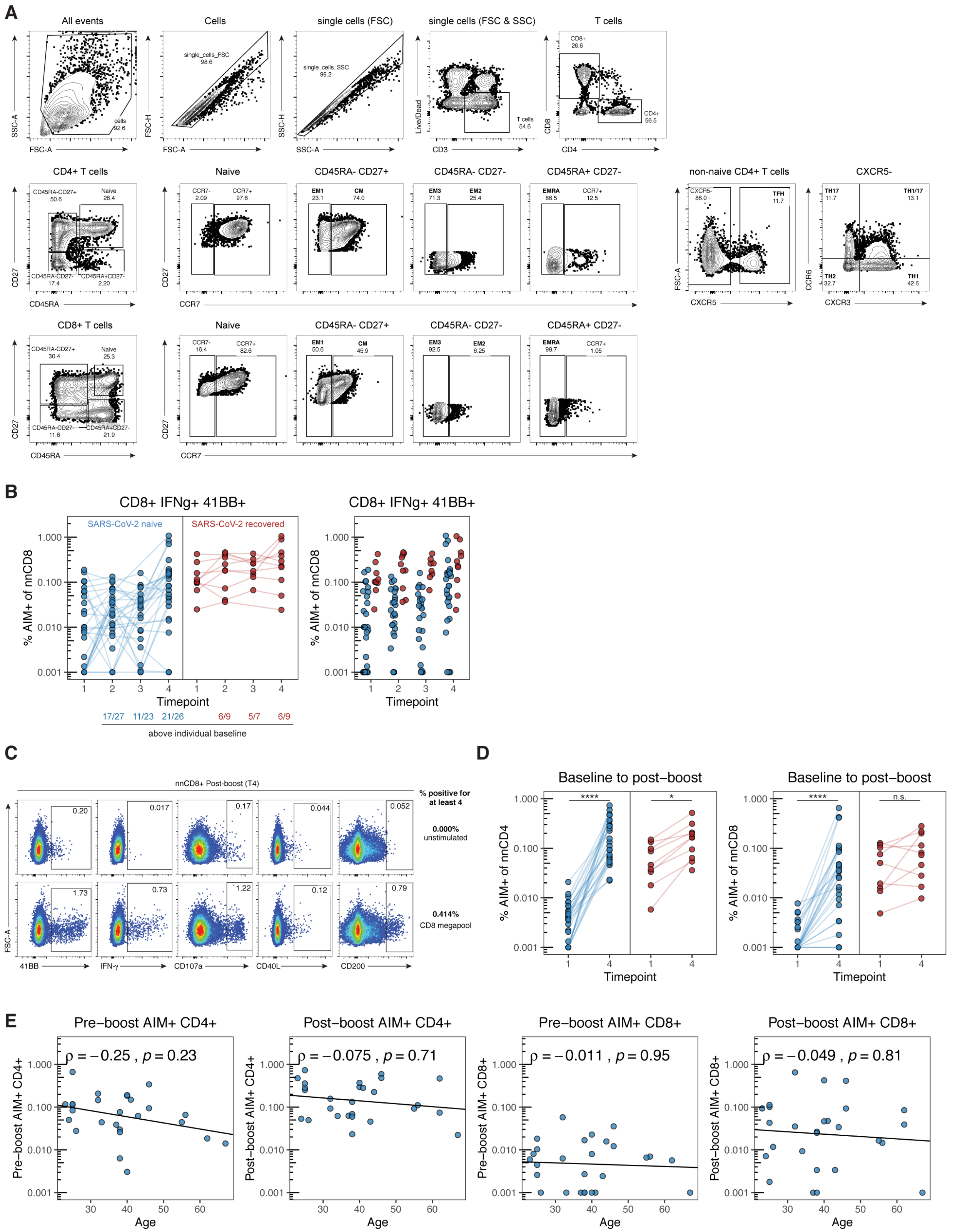


**Fig. S1: Gating strategy and magnitude of AIM⁺ T cell responses.** (A) Gating strategy for identifying T cell subsets. (B) Summary plots of IFN-γ⁺ 41BB⁺ CD8⁺ (right) T cells. Values represent the frequency of IFN-$\gamma^+$ 41BB$^+$ non-naïve CD8$^+$ T cells after subtracting the frequency from paired samples stimulated in the absence of SARS-CoV-2 peptides. Solid lines connect individual donors sampled longitudinally. Statistics were calculated using unpaired Wilcoxon test. (C) Representative flow cytometry plots for identifying AIM$^+$ CD8$^+$ T cells. Numbers represent the frequency of total non-naïve CD8$^+$ T cells. (D) Summary plots of AIM$^+$ CD4$^+$ (left) and CD8$^+$ (right) T cells. Values represent the frequency of AIM$^+$ non-naïve T cells after subtracting the frequency from paired samples stimulated in the absence of SARS-CoV-2 peptides. Solid lines connect individual donors sampled at baseline (timepoint 1) and 1 week post-boost (timepoint 4). Statistics were calculated using unpaired Wilcoxon test. (E) Correlations between the background-subtracted frequency of pre-boost (timepoint 2) or post-boost (timepoint 4) AIM$^+$ CD4$^+$ or AIM$^+$ CD8$^+$ T cells with age. Only SARS-CoV-2 naïve donors were considered for these correlations. Associations were calculated using Spearman rank correlation and are shown with Pearson trend lines for visualization. Blue represents SARS-CoV-2 naïve individuals, red represents SARS-CoV-2 recovered individuals. Timepoints are as defined in Fig. 1A.

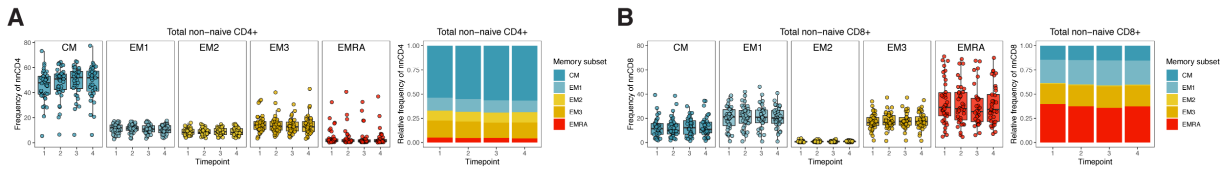


**Fig. S2: Summary of memory T cell subsets in total non-naïve T cells.** (A-B) Frequency of memory T cell subsets in total non-naïve CD4$^+$ (A) and total non-naïve CD8$^+$ (B) T cells. Left panels depict the percent of total non-naïve T cells that are in each subset. Right panels depict the relative frequency of each memory T cell subset in the total non-naive population. CM = CD45RA$^-$ CD27$^+$ CCR7$^+$, EM1 = CD45RA$^-$ CD27$^+$ CCR7$^-$, EM2 = CD45RA$^-$ CD27$^-$ CCR7$^+$, EM3 = CD45RA$^-$ CD27$^-$ CCR7$^-$, EMRA = CD45RA$^+$ CD27$^-$ CCR7$^-$. Timepoints are as defined in Fig. 1A.

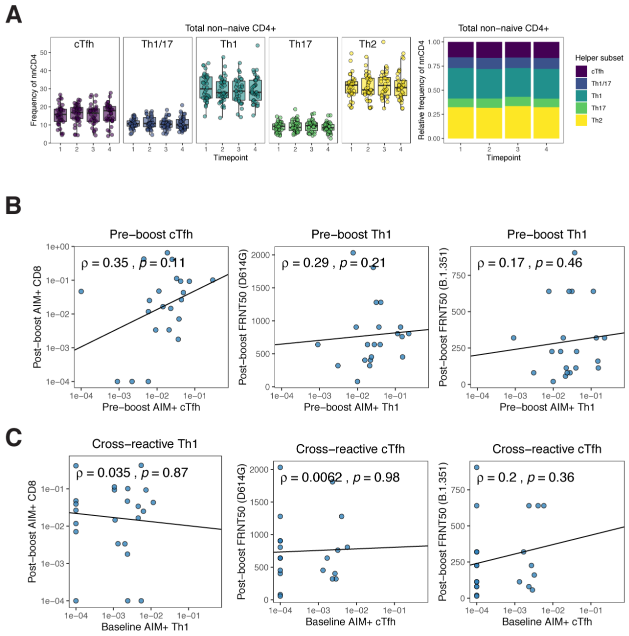


**Fig. S3: CD4$^+$ helper T cell subsets supplement.** (A) Frequency of T helper subsets in total non-naïve CD4$^+$ T cells. Left panel depicts the percent of total non-naïve T cells that are helper T cells in each subset. Right panel depicts the relative frequency of each helper T cell subset in the total non-naive population. cTfh = CXCR5$^+$, Th1 = CXCR5$^-$ CXCR3$^+$ CCR6$^-$, Th2 = CXCR5$^-$ CXCR3$^-$ CCR6$^-$, Th17 = CXCR5$^-$ CXCR3$^-$ CCR6$^+$, Th1/17 = CXCR5$^-$ CXCR3$^+$ CCR6$^+$. (B-C) Correlations between the background-subtracted frequency of pre-boost (B, timepoint 2) and baseline (C, timepoint 1) AIM$^+$ Th1 or AIM$^+$ cTfh cells with post-boost (timepoint 4) AIM$^+$ CD8$^+$ T cells or neutralizing titers against dominant (D614G) or variant (B.1.351) strains of SARS-CoV-2. FRNT50 = Focus reduction neutralization titer 50%. Only SARS-CoV-2 naïve donors were considered for these correlations. Associations were calculated using Spearman rank correlation and are shown with Pearson trend lines for visualization. Timepoints are as defined in Fig. 1A.

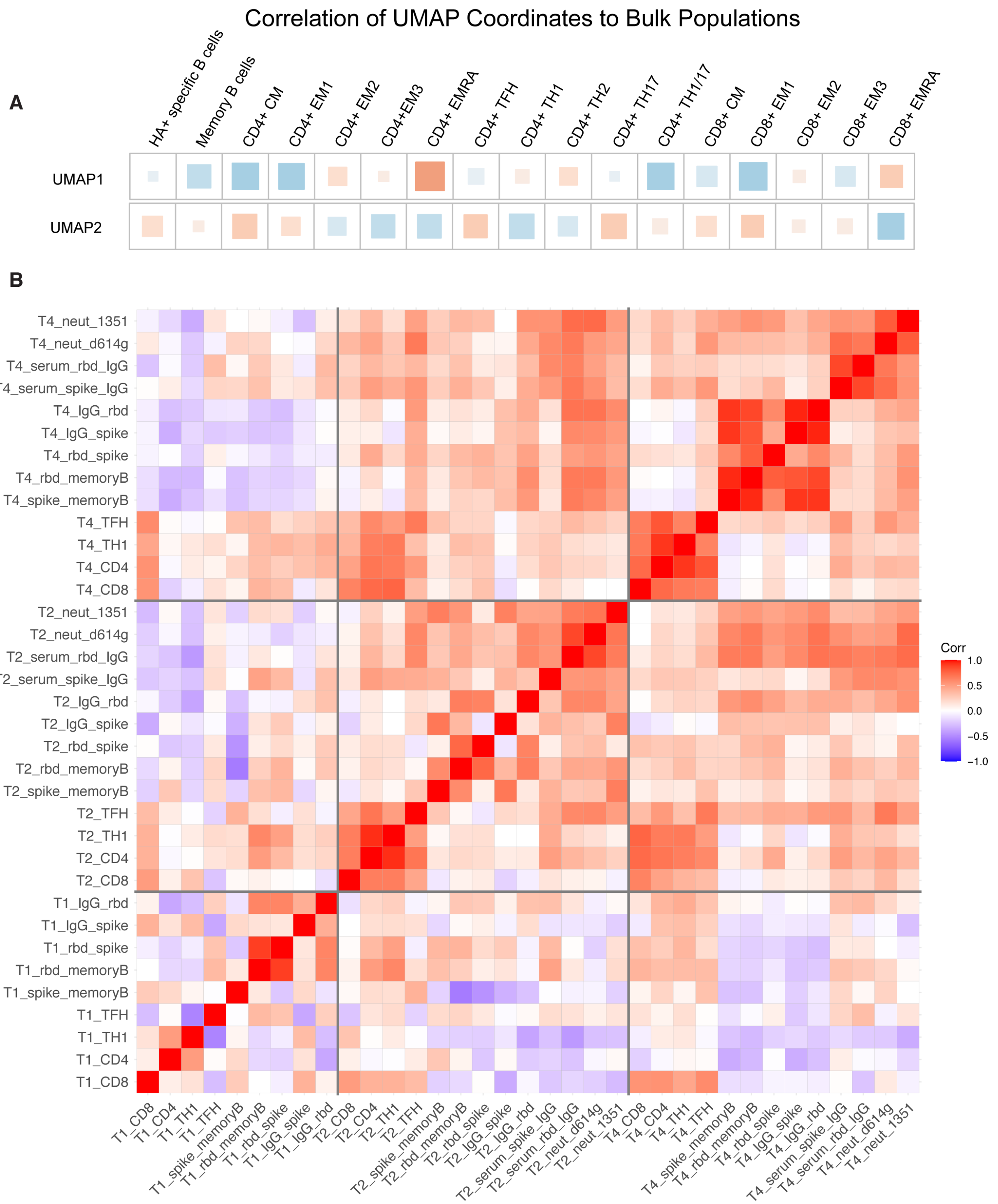


**Fig. S4: Integrated analysis supplement.** (A) Correlations of individual total immune cell populations against the UMAP1 and UMAP2 axis. Red indicates positive correlations and blue indicates negative correlations. (B) Correlations of antigen-specific features over time in SARS-

CoV-2 naïve donors. Associations were calculated using Spearman rank correlation. Timepoints are as defined in Fig. 1A.
